# Isolation, identification and molecular characterization of *Aeromonas* spp. from farmed loach (*Misgurnus anguillicaudatus*)

**DOI:** 10.1101/2021.09.13.460194

**Authors:** Xin Wang, Jiwen Pan, Liqing Chen, Roushan Li, Yu Han, Zihao Di, Bo Ling, Ashfaq Ahmad, Nuo Yang, Lixia Fan, Qian Li, Jifeng Zeng, Guiying Guo, Jiping Zheng

## Abstract

Gram-negative *Aeromonas* bacteria is known to contaminate aquaculture products and retains the ability to infect wide range of host including fishes, shrimps and humans. This study is designed to evaluate the presence of *Aeromonas* in healthy loaches (*Misgurnus anguillicaudatus*) collected from the markets in Hainan and Guangdong provinces of China. Based on the molecular sequencing and phylogenetic analysis with single 16S rRNA and concatenated genes (*gyr*A and *rop*D), 104 isolates were identified as *Aeromonas* species followed by further classification to *A. veronii* (83.65%), *A. jandaei* (9.62%), *A. hydrophila* (3.85%) and *A. allosaccharophila* (2.88%). More than half of the isolates displayed hemolytic activity of 87.5% and were able to form moderate biofilm (78.26%). Fourteen antibiotics from ten representative classes were screened that demonstrated complete resistance to ampicillin, 89.4% and 68.3% to lincomycin and nalidixic acid. Moreover, a notable detection and prevalence was found in screening of ten virulence-related and nine resistance genes. To our knowledge, it is the first report of its kind demonstrating potential threat of the commercial loaches carrying *Aeromonas* to human health. These findings will assist professionals working in clinical settings to optimize their prescription accordingly and researchers to access the impacts of *Aeromonas* resistance on human health.

**Importance:** This study demonstrates high infection rate of loaches by *Aeromonas* spp. in Hainan and Guangdong provinces of China.

The most prevalent species was *Aeromonas veronii*.

Most of the isolates displayed hemolytic activity and formed moderate biofilm.

Multiple virulence genes, resistance genes and multi-antibiotic resistance were detected.

Aeromonads risks should be considered in loach industry.

## 1. Introduction

*Aeromonas* species are Gram-negative pathogenic bacteria, found in aquatic environments like estuary and fresh water. The species of *Aeromonas* can also be obtained from variety of infectious regions of animals, and humans and can cause gastroenteritis, septicemia and wound infections thus presenting a serious threat to human health (Janda and Abbott, 2010; Figueras and Beaz-Hidalgo, 2015; Fernández-Bravo, 2019). The presence in diverse range of infections both in terrestrial and aquatic life, *Aeromonas* spp. have become a fascinating target for research and drug development. Upon infecting fishes, symptoms like bleeding and ulcer are appeared followed by a large-scale deaths and resulting huge economic losses to the aquaculture industry (Janda and Abbott, 2010; Beaz-Hidalgo et al., 2010).

Loach or *Misgurnus anguillicaudatus* is famous for delicious meat and high nutritional value (Zhu and Wu, 2018) that is widely distributed in streams and ditches of Asian countries. Loach is known for high growth rate and for the reason cultivars prefer breeding in large scale (Zhu et al., 2016; Gao et al., 2017). Only in Jiangsu province of China, loach has become a pivotal industry (Wang et al., 2020) yielding annual output of approximately 80 thousand tons of meat and a business of more than one billion yuan. Apart from China, the annual demand for loach in South Korea and Japan is on the rise. In 2014, 90% of loaches in Dunshang county of Jiangsu province was exported to South Korea and Japan (Zhang et al., 2020). Infections in loach not only lead to economic losses but also reduces the quality of food and posing health threats to humans (Wang et al., 2020). For instance, *Aeromonas veronii* has caused severe harm to loach industry, which can cause ulcerative syndrome or skin congestion (Jun et al., 2010; Wang et al., 2020).

Since 1943, total of 36 species of *Aeromonas* have been described which can further be classified into motile mesophilic and non-motile psychrophilic strains. The mesopphiles, like *A. hydrophila*, causes human infections while the psychrophiles, like *A. salmonicida*, infect fishes. Due to similar phenotypes, *Aeromonas* spp. pose difficulty in identification and could lead to misjudgment (Beaz-Hidalgo et al., 2010). For instance, *A. dhakensis* was initially identified as *A. hydrophila* and was reclassified in 2012 (Beaz-Hidalgo et al., 2013). Due to higher degree of similarity in *Aeromonas* spp. the universal marker of 16S rRNA may generate false negative results therefore, the combination of housekeeping genes like *gyr*A and *rpo*D are found helpful in identification of new species. (Martinez-Murcia et al., 2011; Khor et al., 2015; Miñana-Galbis et al., 2013; Chun and Bae, 2000).

The pathogenicity of *Aeromonas* spp. is associated to virulence factors like extracellular proteins (cytotoxic enterotoxin, hemolysin, cytotoxin), structural components (flagella), secretion systems and quorum sensing (Fernández-Bravo and Figueras, 2020). For the purpose of addressing *Aeromonas* pathogenicity, antibiotics are directly poured into the aquaculture that somehow assist bacteria to evolve and generate resistance (Guz and Kozińska, 2004). Recent published reports have detected highly resistant *A. veronii* strains and multidrug-resistant *Aeromonas* spp. in clinical and environmental samples (Dias et al., 2012; Yano et al., 2015).

This study design investigated the prevalence of *Aeromonas* spp. in loaches of Guangdong and Hainan provinces of China. Besides, we evaluated the diversity of *Aeromonas* species using housekeeping genes as reference markers, the capacity to form biofilm, the presence of virulence-related and drug resistance genes.

## 2. Materials and methods

### 2.1. Sampling of loach and isolation of *Aeromonas* spp

From 2019 to 2020, total of 152 loach samples were collected in five batches from random farms in Hainan and Guangdong provinces of China. Among them, four batches were collected in 2019 and one batch was collected in 2020. All the samples were dissected in a sterile environment, followed by isolation and purification from the lesions of gill, skin, and viscera. All cells were inoculated on LB solid medium containing 50 μg/ml ampicillin and cultured overnight at 30°C. Strains showed prominent growth were selected and sub-cultured on LB solid medium containing 50 μg/ml ampicillin overnight at 30°C. The collected strains were transferred to LB liquid medium containing 50 μg/ml ampicillin and cultured overnight at 30°C to obtain pure culture strain. Finally, the strain was stored in the mixture of liquid medium: 30% glycerol (1:1) at - 80°C until identification is completed.

### 2.2. Identification of isolates

Gram staining, physiological and biochemical kits were used to identify the colonies, followed by the use of 16S rRNA, *gyr*A and *rpo*D housekeeping gene for molecular identification. To observe the growth state, the purified strains were inoculated on LB solid medium retaining 50 μg/ml ampicillin at 30 °C for 24 h. A single colony was selected for Gram staining, whereas morphology and size were observed under the microscope. The purified strains were re-inoculated and conventional methods like “Identification Manual of common bacteria” were applied for biochemical characteristics. Next, genomic DNA of the isolates were extracted using genomic DNA Extraction Kit (Nanjing novozan Biotechnology Co., Ltd., China). Using primers, the 16S rRNA, *gyr*A and *rpo*D housekeeping genes were amplified through PCR employing 2 × A8 Taq PCR master mix (Beijing Adlay Biotechnology Co., Ltd., China). The reaction procedures were as follows: 94°C for 3 min, 94°C for 30 s, 54°C for 30 s, 72°C for 1 min for total of 30 cycles, 72°C for 10 min. PCR products were sequenced commercially at Guangzhou Tianyi Huiyuan Biotechnology Co., Ltd., China. For analysis of the sequenced data, ClustalW and MEGA X software were utilized with 1000 bootstraps. Phylogenetic analyses were carried out by neighbor-joining method (Thompson et al., 1994) and the final tree was prepared with reference sequences (Letunic and Bork, 2007). All the reference sequences were obtained from the NCBI (Supplementary Table 1).

### 2.3. Molecular characteristics of the potential virulence factors

The hemolytic activity of 104 *Aeromonas* spp. were determined on 5% (v/v) sheep blood agar plate (Guangdong huankai Microbial Technology Co., Ltd., China). In case of clear area beside the line, the hemolytic test was considered positive. We used the already reported PCR primers in Supplementary table 2 that detected ten ten virulence factor genes including the hemolysin (*hyl*A), cytotoxic enterotoxin (*act*), cytotoxic enterotoxins (*alt, ast*), flagella (*fla*), lipase (*lip*), elastase (*ela*) and three of the secretory system T3SS (*asc*V and *aex*T), and aerolysin (*aer*), respectively. PCR amplification was performed in final volume of 25 μL, including 12.5 μl of 2 × F8 Taq PCR master mix (Nanjing novozan Biotechnology Co., Ltd., China), 1 μl of 10 μm primers, 1 μl of DNA template (30-40 ng) and 10.5 μl of ddH_2_O. The cycle conditions include initial single cycle at 94°C for 3 minutes, followed by 30 cycles, including denaturation at 94 °C for 10 seconds, annealing at 47°C - 56°C for 15 seconds, elongation at 72°C for 10 seconds, and finally a cycle of 5 minutes at 72°C. PCR results were confirmed by 0.8% agarose gel electrophoresis.

### 2.4. Detection of antimicrobial susceptibility and resistance genes

According to the current guidelines of the Clinical and Laboratory Standard Institute (CLSI, 2012), Kirby-Bauer (K-B) disk diffusion method was used for the drug sensitivity test. The sensitivity of *Aeromonas* spp. against 14 antibiotics was tested (Table 2). All the strains were divided into highly sensitive, moderately sensitive, and drug-resistant according to the CLSI and manufacturer standards. *Escherichia coli* (ATCC25922) was used as the quality control strain for the drug sensitivity test.

Eleven resistance genes were detected using conventional PCR. The primers details are shown in Supplementary table 2. These resistance genes include β-lactam, *tem, ctx*M, *cph*A; tetracyclines, *tet*A, *tet*C; sulfanilamide: *sul*1; quinolones: *qnr*A, *qnr*S; linxamines: *lin*A*/lin*A*’*; amido alcohols: *cat*A2; and macrocyclic lipids: *erm*B were detected. PCR amplification was performed at a final volume of 25 μl, including 12.5 μl of 2 × F8 Taq PCR mastermix, 1 μl of 10 μm primers, 1 μl of DNA template (30-40 ng) and 10.5 μl of ddH_2_O. The cycle conditions include initial single cycle at 94°C for 5 minutes, followed by 30 cycles, including denaturation at 94°C for 30 seconds, annealing at 47°C - 56°C for 30 seconds, elongation at 72°C for 30 seconds, and finally a cycle of 5 minutes at 72°C. Gel electrophoresis was used as above to confirm PCR product.

### 2.5. Biofilm formation

To analyze capacity of biofilm formation, *Aeromonas* strains were cultured overnight at 30°C on LB solid medium. Single colonies were selected and inoculated in fresh LB liquid medium at 30°C and kept shaking on 180 rpm for 16 h. The OD_590_ values of the two bacterial solutions were adjusted to 1.0 by spectrophotometer, diluted into the fresh LB liquid medium at the ratio of 1:40, and 200 μl each was added into 96 well plate. Three replicates were set in each group and incubated at 30°C for 24 h. The liquids were aspirated followed by the addition of add 200 μl sterile PBS buffer to each well. Rinsing was performed in triplicate and turn it upside down to dry. 250 μl methanol was added to each well for 15 min, and discarded for an air dry. Next, 100 μl of crystal violet solution (0.1%) was added to each well and dye them for 5 min. With the help of sterile PBS, staining solution was aspirated and made them dry. 100 μl of 95% ethanol was added to each well, and the biofilm was dissolved at 30°C for 30 min. OD_590_ value was determined by microplate reader.

## 3. Result and Discussion

### 3.1. Identification of *Aeromonas* species

Total of 104 samples were collected and purified from the lesions of gills, skin, and viscera of five different batches of loach (152 loaches). Microscopic analyses revealed the morphology of short, small and mostly found in a single arrangement. Based on Gram staining, biochemical and physiological tests, all the 104 isolates were identified as *Aeromonas* spp. and were found oxidase-positive, that may retain the ability to ferment glucose and mannitol but may not ferment inositol. The detection analyses based on the concatenated sequenced data of *gyr*A and *rpo*D against the reference sequences highlighted the presence of four *Aeromonas* species. Among them 87 strains (83.65%) of *A. veronii*, 10 strains (9.62%) of *A. jandaei*, 4 strains (3.85%) of *A. hydrophila* and 3 strains (2.88%) of *A. allosaccharophila*, were classified (Figure 1). Relating to the geographical location, of 87 strains of *A. veronii* 26 and 61 were obtained from Hainan and Guangdong provinces whereas all the identified 3 strains of *A. allosaccharophila* were collected from Guangdong province. Likewise, 7 and 1 strains of *A. jandaei* and *A. hydrophila*, turned out from Hainan while and 3 each from Guangdong province (Supplementary Table 3).

**Figure 1.**
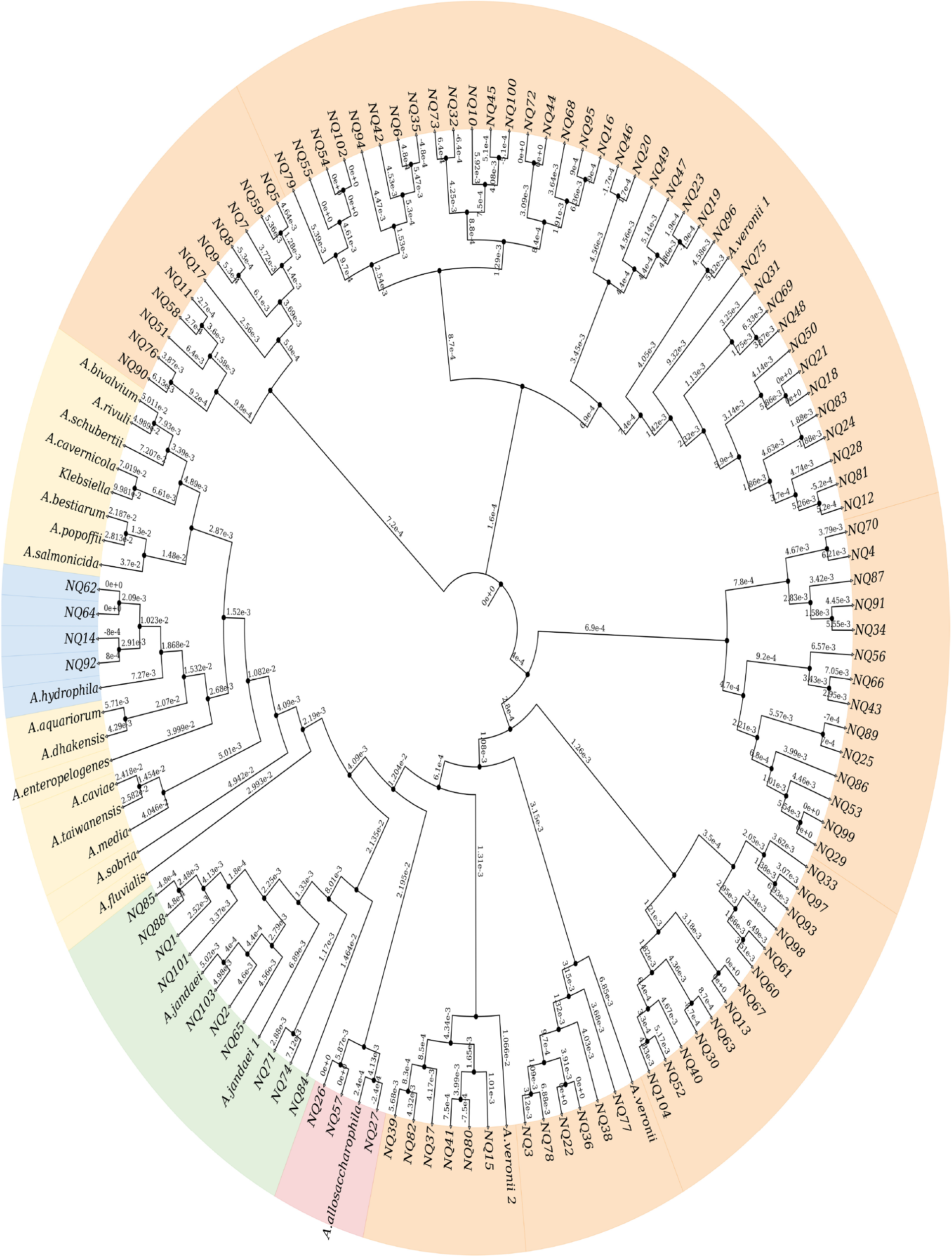
Phylogenetic tree based on *gyr*A and *rpo*D concatenated housekeeping gene sequences showing a relationship within the genus *Aeromonas* of 104 isolates from loach collected from Guangdong and Hainan. The tree was constructed by the neighbor-joining method with Kimura’s two parameter method. Bootstrap values based on 1000 replicates are shown. The areas of orange, blue, green and maroon indicate the isolates belonging to *A. veronii, A. hydrophila, A. jandaei* and *A. allosaccharophilla*, respectively, and yellow are the typical strains of the other *Aeromonas* species. (Line 191)

Species of *Aeromonas* are known and as common pathogen in aquatic environment often leading to death. Since 1994, *A. veronii, A. hydrophila*, and *A. jandaei* have been considered the primary or secondary pathogens of fish diseases. *A. veronii* is associated with the large-scale death of variety fishes like Thai catfish, loach and tilapia (Hoai et al., 2019). It is noteworthy to mention that these fish are widely distributed worldwide that itself explain the severe nature of *Aeromonas* infection to the global aquaculture industry. Our results indicated 84% occurrence of *A. veronii* suggesting a potential risk to loach cultures in Hainan and Guangdong provinces. Besides, *A. veronii, A. hydrophila*, and *A. dhakensis* are the most common isolated in the clinical settings (Figueras and Beaz-Hidalgo, 2015). *A. hydrophila* appeared for the first time in 2006 followed by the classification of *A. dhakensis* which is common to coastal regions of tropical and subtropical climates (Seshadri et al., 2006). On toxicity scale, the higher toxicity of *A. dhakensis* than *A. veronii, A. hydrophila* and *A. caviae* has been reported previously (Chen et al., 2016), however, in the current study we did not detect the presence of *A. dhakensis* in loach samples but found it in the farmed crocodiles with pneumonia and septicemia in Hainan (Pu et al., 2019). Recently, Zhu and coworkers have isolated *A. veronii* from the skin of ulcerated loaches of Jiangsu province (Zhu et al., 2016). Regarding the fact that loach is a popular and food of choice thus favors the transmission route of *Aeromonas* spp. to humans. We have isolated all of our samples from the loaches available in the market to the consumer indicates that these loaches have been infected before they reach to the market. The *Aeromonas* spp. isolated from the skin and viscera of loaches may potentially cause systemic infections and therefore, a proper heat treatment during processing is necessary to achieve the bactericidal effect.

### 3.2. Distribution of virulence genes

It has been mentioned that hemolysins and extracellular hydrolytic enzymes of *Aeromonas* spp. assist invasiveness and establishment of infections (Chacón et al., 2003). To access the hemolytic nature of the isolated strain, we examined hemolytic activity of 104 *Aeromonas* isolates where 91 (87.5%) isolates displayed hemolytic activity. Next, the virulence genes of *Aeromonas* spp. were screened that included the TTSS genes (*asc*V and *aex*T), the enterotoxin and hemolysin genes and lipase, elastase, and flagella genes (*lip, ela*, and *fla*) (Table 1). Total of 70 (67.3%) and 77 (74%) isolates tested positive for *asc*V and *aex*T genes. The ascV and aexT genes were found in all *A. allosaccharophila* isolates whereas 66.7% and 79.3% were detected in A. veronii, 50% each in *A. hydrophila*, 70% and 30% in *A. jandaei*, respectively.

**Table 1.** Virulence factor genes of *Aeromonas* spp. isolates from loach.

| Species |  | Number and (%) of positive isolates |  |  |  |  |  |  |  |  |  |
| --- | --- | --- | --- | --- | --- | --- | --- | --- | --- | --- | --- |
|  |  | *hylA* | *ascV* | *aexT* | *aer* | *act* | *ast* | *alt* | *lip* | *ela* | *fla* |
| *A.veronii* | (n=87) | 56 (64.4) | 58 (66.7) | 69 (79.3) | 87 (100) | 86 (98.9) | 18 (20.7) | 37 (42.5) | 52 (59.8) | 55 (63.2) | 80 (92.0) |
| *A.allosaccharophila* | (n=3) | 3 (100) | 3 (100) | 3 (100) | 3 (100) | 3 (100) | 0 | 1 (33.3) | 2 (66.6) | 3 (100) | 2 (66.6) |
| *A.hydrophila* | (n=4) | 3 (75) | 2 (50) | 2 (50) | 4 (100) | 4 (100) | 1 (25) | 3 (75) | 1 (25) | 4 (100) | 4 (100) |
| *A.jandaei* | (n=10) | 5 (50) | 7 (70) | 3 (30) | 10 (100) | 9 (90) | 1 (10) | 2 (20) | 3 (30) | 6 (60) | 6 (60) |
| Total | (n=104) | 67 (64.4) | 70 (67.3) | 77 (74.0) | 104 (100) | 102 (98.1) | 20 (19.2) | 43 (41.3) | 58 (55.8) | 68 (65.4) | 92 (88.5) |

*hylA* (haemolysin), *ascV* and *aexT* (type three secretion systems), *aer* (aerolysin), *act* (cytotoxic enterotoxin), *ast* and *alt* (cytotoxic enterotoxins), *lip* (lipase), *ela* (elastase), *fla* (flagella).

Together the enterotoxin and hemolysin genes (*hyl*A, *aer, act, ast* and *alt*) were turned out positive in 67 (64.4%), 104 (100%), 102 (98.1%), 20 (19.2%) and 43 (41.3%) isolates. Specifically, the *hyl*A gene was detected 100% in *A. allosaccharophila* and 50% to 75% in three *Aeromonas* species. The *aer* gene marked 100% detection in isolated population and likewise the presence of *act* gene stayed high 90%-100%. Contrary, the *ast* gene identified on a lower side having a detection rate of 19.2%. The *alt* gene showed somehow a scattered rate of detection and was found in 42.5% of *A. veronii*, 75% of *A. hydrophila*, 20% of *A. jandaei*, and 66.6% of *A. allosaccharophila*. Lipase, elastase, and flagella genes (*lip, ela*, and *fla*) were detected in 58 (55.8%), 68 (65.4%) and 92 (88.5%) in all 104 isolates. The *lip* gene was found in 59.8% of *A. veronii*, 66.6% of *A. allosaccharophila*, 25% of *A*.*hydrophila*, and 30% of *A*.*jandaei* likewisethe *ela* and *fla* genes were detected in the range of 60% to 100%.

Species of *Aeromonas* are known to secrete enterotoxin and hemolysin which are the main cause of pathogenicity. These results are in agreement with published reports indicated the detection of such virulence genes with different frequencies in *Aeromonas* spp. (Wu et al., 2019; Rasmussen-Ivey et al., 2016). Walsh and coworkers associated the virulence of *Aeromonas* spp. to the cumulative effect of multiple virulent genes (Walsh et al., 1995). This study reports the presence of four genes on average per isolate depicting the severity and potential risk to human health. To date, *Aeromonas* spp. are recognized for six secretion systems involved in virulence factors transportation (Fernández-Bravo and Figueras, 2020), for instance, T3SS injects the effector proteins into the host bacteria and regulate pathogenicity (Chandrarathna et al., 2018). The mutant T3SS in *A. salmonida* and *A. hydrophila* showed reduced pathogenicity than to the wild types further substantiates their importance (Fernández-Bravo and Figueras, 2020). Among the others, the *fla, ela* and *lip* genes assist bacterial colonization, pathogenesis through gene disruption, and nutritional uptake in *Aeromonas* species (Fernández-Bravo and Figueras, 2020).

Overall, we found that these virulence genes are highly heterogeneous and showed differences among species of *Aeromonas*. To compare our detection frequency of three virulent genes, *act, alt* and *ast* with reported from the Asian countries, we found 98.1%, 41.3%, and 19.2% compared to 100%, 39-42%, and 0-8% reported (Yano et al., 2015; Puthucheary et al., 2012; Yi et al., 2013). Since the importance of loach as direct human food, further work is needed to elucidate the impacts on human health.

### 3.3. Antibiotic sensitivity

Bacteria retain the ability of evolving to equip and resist variety of threats in the form of antibiotics or stress (Ahmad et al., 2020). To understand the nature of antibiotic sensitivity among these samples, the disk diffusion method was employed for fourteen commonly used antibiotics. These fourteen candidates were selected from the antibiotic classes of aminoglycoside, cephalosporins, carbapenem, monobactam, macrolide, tetracycline, lincosamide, sulfonamide, fluoroquinolone and quinolone. All the identified species were found resistant to ampicillin followed by high ranges of lincomycin and nalidixic acid where the resistance frequency was up to 82% and 68.3%. Towards, cefotaxime, gentamicin and chloramphenicol these isolates showed considerably lower resistance, i.e., <12%. Regarding cephalotin, complete resistance was found in *A. hydrophila* and *A*. jandaei, however 88% and 33% sensitive profile was recorded for *A. veronii* and *A. allosaccharophila*. The resistance rates to ampicillin-sulbactam, tetracycline, norfloxacin, trimethoprim-sulfamethoxazole, imipenem and erythromycin were found in moderate range of 46.2%, 28.8%, 32.7%, 28.8%, 49.0 %, 47.1%, respectively (Table 2). All the isolates were found sensitive to aztreonam and in totality 89.4% of the species were found resistant to three and more antibiotics suggesting low sensitivity or higher resistance.

**Table 2.**
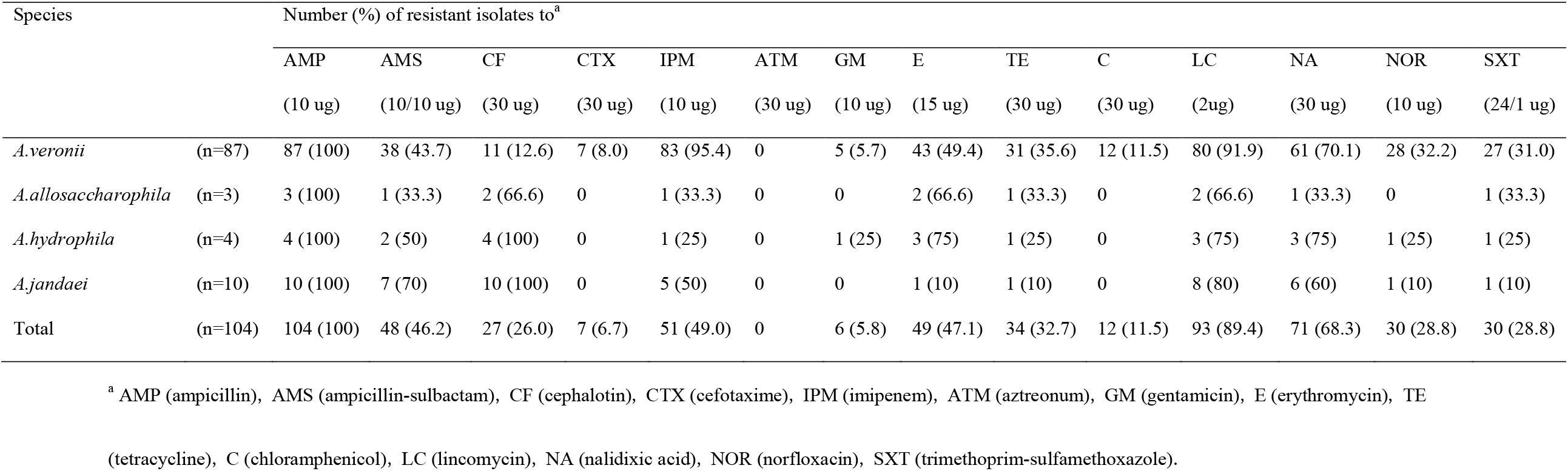
Resistance to antimicrobial agents of *Aeromona*s spp. isolates from loach.

Next, we opted to screen the drug-resistant genes mentioned in table 3. The screening results identified metalo-β-lactamase genes entailing *tem, ctx*M and *cph*A in 60.6%, 12.5% and 98.1% of the isolates. The *tem* gene displayed complete detection in *A*.*hydrophila* while 33.3% each in *A. allosaccharophila* and *A. jandaei* and 58.6% in *A. veronii*. Contrary to the *tem* gene, *ctx*M was detected in 13 samples of *A. veronii* though *cph*A was rendered at a high rate of 98.1%. Among other tested genes, the tetracycline resistant genes, *tet*A, *tet*C, the linexamines genes, *lin*A*/lin*A, the amido alcohols gene, *cat*A2, and the macrolides *ermB* exhibited lower or no detection in the tested four species (104 isolates). The detection rate of sulfonamides resistant genes *Sul1* appeared on the higher side, attaining 77.9% in all isolates whereas quinolones *qnr*A and *qnr*S displayed 51.9% and 42.3%, respectively.

**Table 3.** Antibiotic resistant genes of *Aeromonas* spp. isolates from loach.

| Species |  | Number and (%) of positive isolates |  |  |  |  |  |  |  |  |  |  |
| --- | --- | --- | --- | --- | --- | --- | --- | --- | --- | --- | --- | --- |
|  |  | *tem* | *ctxM* | *cphA* | *tetA* | *tetC* | *sul1* | *qnrA* | *qnrS* | *linA/linA'* | *catA2* | *ermB* |
| *A.veronii* | (n=87) | 51 (58.6) | 13 (14.9) | 85 (97.7) | 25 (28.7) | 21 (24.1) | 70 (80.1) | 48 (55.2) | 33 (37.9) | 0 | 0 | 0 |
| *A.allosaccharophila* | (n=3) | 1 (33.3) | 0 | 3 (100) | 1 (33.3) | 2 (66.6) | 2 (66.6) | 2 (66.6) | 3 (100) | 0 | 0 | 0 |
| *A.hydrophila* | (n=4) | 3 (100) | 0 | 4 (100) | 0 | 1 (50) | 3 (75) | 2 (50) | 3 (75) | 0 | 1 (25) | 0 |
| *A.jandaei* | (n=10) | 8 (33.3) | 0 | 10 (100) | 0 | 2 (20) | 6 (60) | 2 (20) | 5 (50) | 0 | 0 | 0 |
| Total | (n=104) | 63 (60.6) | 13 (12.5) | 102 (98.1) | 26 (25) | 26 (25) | 81 (77.9) | 54 (51.9) | 44 (42.3) | 0 | 1 (0.96) | 0 |

The resistance of *A. veronii* to variety of antibiotics has been reported (Yang et al., 2017). Particular to *A. veronii*, the higher resistance was displayed to aminoglycoside, carbapenem, licosamide, macrolid and quinolone class of antibiotics (Figure 2). Similar results are reported for catfish in Vietnam where high resistance was observed for carbapenem, lincosamide and macrolide (Hoai et al., 2019). To treat loach-derived infection of *Aeromonas* spp., Yu and coworkers have suggested the use of gentamicin (aminoglycoside) or erythromycin (macrolide) in South Korea, and only gentamicin in Guangdong and Hainan provinces of China (Yu et al., 2015). Previously, *A. hydrophila* was found in complete resistance to ampicillin and cephalotin (Goni-Urriza et al., 2000). In the case of carbapenem resistance we observed inconsistent data, i.e., 50 and 54 of the total isolates were collected in four bathes in the year 2019 and 2020. The batches collected in 2019 and 2020 have displayed ∼82% and ∼18.5% of resistance to imipenem suggesting drastic changes to carbapenem. Previous reports concluded that the excessive use of cefotaxime and imipenem in clinical settings against *Aeromonas* spp. favor antibiotic resistance in patients and thus necessary to use them with care (Sen and Rodgers, 2004; Wu et al., 2011; Chen et al., 2014).

**Figure 2.**
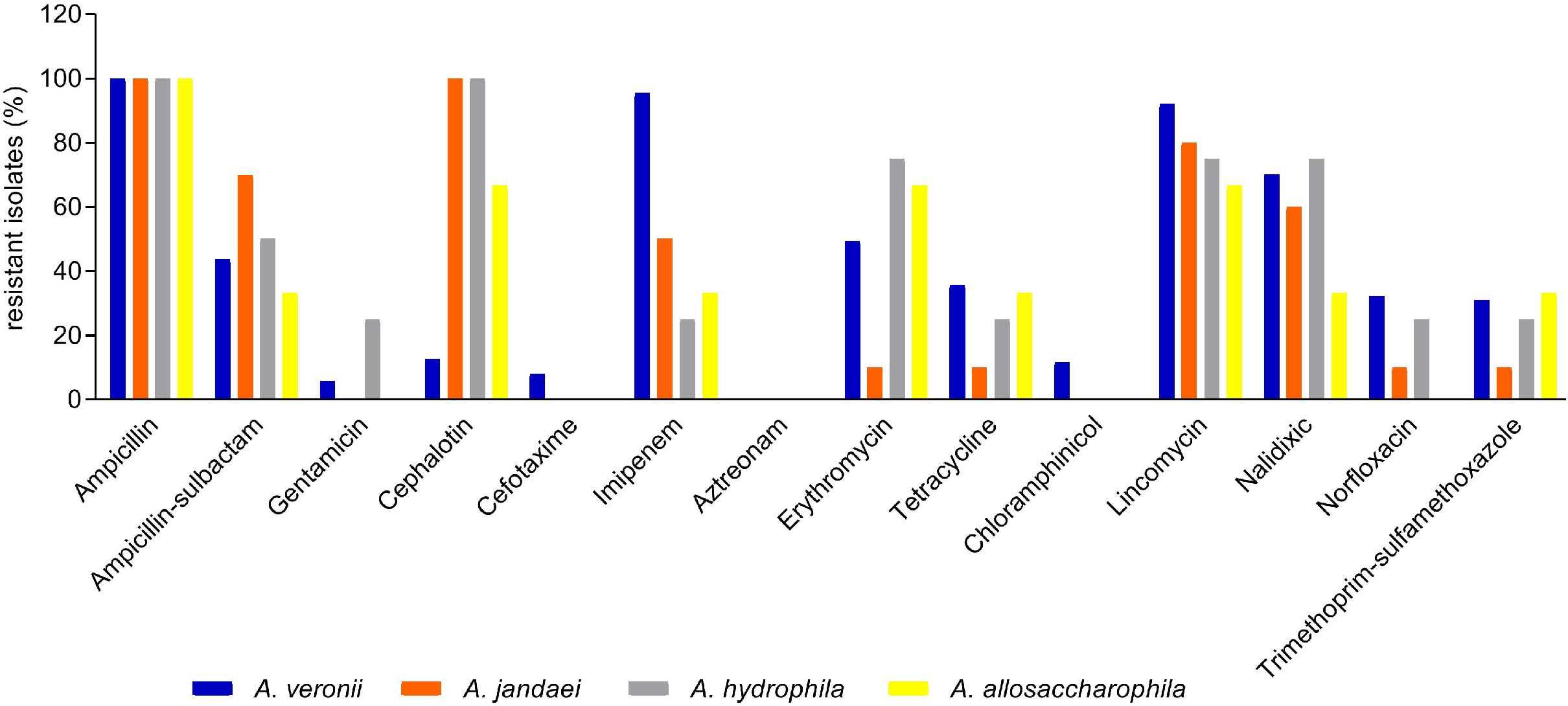
Resistance profile of Aeromonas species to various classes of antibiotics. Antibiotics candidates from ten classes were tested against *A. veronii* (blue), *A. jandaei* (orange), *A. hydrophila* (grey) and *A. allosaccharophila* (yellow). Vertical bars indicate species resistance in percent whereas absence of bar denotes complete sensitivity. (Line 291)

To this side, we detected number of drug resistance genes in these *Aeromonas* strains but remained unable to classify them with particular resistance and further study is required to explore their direct or indirect involvement in resistance. Our findings suggest that *Aeromonas* spp. are resistant to variety of antibiotics in aquaculture, therefore, precautionary strategies are needed in selection of antibiotics for treatment in loach culture. To avoid bio-accumulation it is necessary to limit the use of such antibiotics that have already developed resistance.

### 3.4. Quantitative analysis of biofilm

Bacteria retain the ability to form biofilms, assisting them in anchoring and enable increased resistance to antimicrobial agents and pathogenesis (Ge et al., 2018). To identify the ability of biofilm formation, the OD_590_ values were measured by the dissolved crystal violet and compared against the negative control (LB medium). Our result indicated that majority of the species are able to form moderate biofilm (n=18,78.26%). Apart from one isolate (4.35%) the remaining 17.39% were proficient to form weak biofilm (Table 4).

**Table 4.**
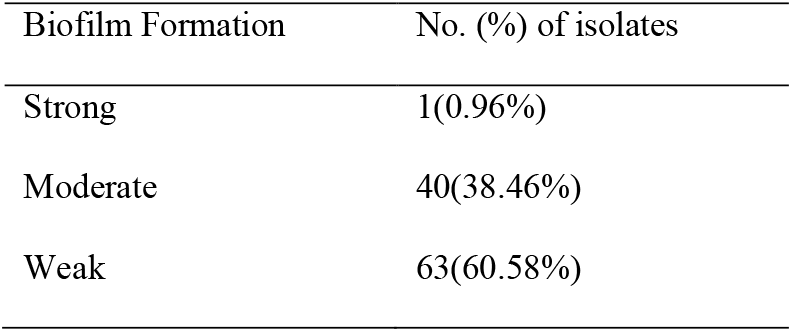
Detection of biofilm.

*Aeromonas* spp. require biofilm to attain the attachment and fulfill nutritional needs. These species are able to gain environmental adaptation against salinity and antibiotics and to enhance virulance through alteration in biofilm (Talagrand-Reboul et al., 2017; Odling-Smee et al., 2003; Peterson et al., 2015; Angeles-Morales et al., 2012; Van Acker et al., 2014; Zanella et al., 2012). On the other side, aquaculture provide intensive feeding mode that contain high content of organic matter in large scale may lead to the outbreak of disease and recurrence infection (Basson et al., 2008). We observed the formation of moderate biofilm associated with high virulence and drug resistance. Loach lives in shallow and silt containing water retaining high concentration of antibiotics that basis for strong resistance in the isolated *Aeromonas* spp.

## 4. Conclusion

In conclusion, this study reports high infection rate of loaches by *Aeromonas* spp. in Hainan and Guangdong provinces of China. The presence of virulent genes, biofilm formation and the drug resistance profile suggest that loach aquaculture industry should pay attention to the classes and concentration of antibiotics in future breeding. The detected resistance profile to all the major classes of antibiotics is an alarming one. In order to minimize the risk of *Aeromonas* infection in humans, hygienic principles should be followed during cooking.

## Declaration of competing interest

The authors declare that this manuscript has not been submitted to, nor is under review at, another journal or other publishing venue. None authors have affiliation with any organization with a direct or indirect financial interest in the subject matter discussed in the manuscript.

## Acknowledgments

This work was supported by the Natural Science Foundation of China (32060788 to J. Zeng and 32060131 to J. Zheng) and the Basic and Applied Basic Research (natural Science) of Hainan Province for High-level Talents program (Haikou Bureau of Science and Technology) (2019RC084 to J. Zheng).

## Authors contribution

Jiping Zheng, Jifeng Zeng and Guiying Guo conceived the experiments. Xin Wang, Jiwen Pan, Liqing Chen, Roushan Li, Yu Han, Zihao Di, Bo Ling, Nuo Yang, and Lixia Fan performed the experiments. Jifeng Zeng, Xin Wang and Qian Li analyzed the results. Xin Wang, Ashfaq Ahmad and Jiping Zheng wrote the paper. All authors read and approved the final manuscript.

## Availability of data and materials

Additional Supporting Information may be found in the online version of this article:

Supplementary Table 1. Reference sequences of *Aeromonas*

Supplementary Table 2. Details of the oligonucleotide primers used in the study

Supplementary Table 3. Distribution and sampling of *Aeromonas* in loach the 104 *Aeromonas* strains and the gene sequences of *gyr*A and *rpo*D were accessible via Genbank website (https://www.ncbi.nlm.nih.gov/genbank/). The accession numbers were: MZ494451 to MZ494457 and MZ643467 to MZ643563 for *gyr*A and MZ643564 to MZ643667 for *rpo*D.

